# Protective Effects of Butyrate on Retinal Neovascularization in Preclinical Retinopathy of Prematurity Models

**DOI:** 10.1101/2024.06.03.597234

**Authors:** Allston Oxenrider, Tommy Bui, John Lester, Gayatri Seth, Aleah J. Brokemond, Menaka C. Thounaojam, Pamela M. Martin, Ravirajsinh N. Jadeja

## Abstract

Retinopathy of prematurity (ROP) remains a leading cause of childhood blindness worldwide, necessitating new therapeutic strategies. Current interventions targeting advanced disease stages often fail to prevent long-term visual impairment. This study investigates the potential of sodium butyrate (NaB), an orally administered short-chain fatty acid, in preclinical models of ROP. Using the oxygen-induced retinopathy (OIR) mouse model, we demonstrate that daily oral NaB supplementation significantly protects against pathological angiogenesis, impacting not only vascular but also neuronal and microglial pathology in the inner retina. Notably, NaB shows efficacy in early-phase ROP intervention, as evidenced by studies in postnatal day 9 (P9) OIR mice and a novel hyperglycemia-associated retinopathy (HAR) model that mimics the hyperglycemic conditions of many premature infants. These findings highlight NaB as a promising alternative or adjunct therapy to current anti-VEGF treatments, offering protection across multiple retinal cell types and stages of ROP development. The study underscores the need for further research to elucidate the specific mechanisms of NaB’s action, paving the way for its potential clinical application in ROP management. This research marks the first exploration of butyrate as a preventative and therapeutic agent for ROP, setting the stage for additional preclinical evaluations and optimization.

## 1. Introduction

Retinopathy of prematurity (ROP) is a leading cause of childhood blindness ^1–3^. Low birth weight (less than 1250 grams), poor postnatal weight gain, low gestational age (less than 31 weeks), and prolonged oxygen therapy have been consistently linked to increased disease susceptibility ^1–3^. It is often difficult to prevent premature delivery and thereby impact the first two of the above listed susceptibility factors. Therefore, it is not surprising that clinical efforts to alleviate the risk of ROP development have focused heavily on tightly regulating oxygen therapy to support proper tissue oxygenation in the absence of complete lung development while limiting unwanted effects. However, even with this, millions of premature babies continue to develop ROP. Thus, it is clear that additional causative factors may be involved at the molecular level. There are no strategies to prevent ROP or intervene early in its development. Existing strategies essentially involve ablation of the peripheral retina through surgical procedures such as laser photocoagulation, cryotherapy, and vitrectomy ^4–6^. Intravitreal injection of anti-vascular endothelial growth factor therapies represents an additional highly utilized option. However, these strategies are capable of impacting only advanced or proliferative (phase II) ROP and have significant associated risks. Atop being highly invasive, current strategies for clinical management of ROP additionally pose the risk of infection or disruption of normal developmental processes such as normal vascular development still taking place in the premature retina. Further, even when the above measures are effective, affected children may still experience vision loss and are left with an increased propensity to develop eye problems later in life, such as myopia, strabismus, and amblyopia ^7–10^. There is a tremendous need for improved therapies to prevent and treat ROP.

With the goal of testing and developing a novel therapy for ROP, the current study focuses on short-chain fatty acid (SCFA) butyrate. SCFAs are secondary metabolites produced from the fermentation of non-digestible carbohydrates by the gut microbiota. SCFAs have diverse regulatory functions and impact host physiology and immunity ^11^. In the infant intestine, the SCFA butyrate is produced by the bacterial fermentation of human milk oligosaccharides ^12^. However, premature birth impacts proper microbiome acquisition and, in turn, overall SCFA production, including that of butyrate ^13–15^. Most premature infants receive antibiotic therapy to prevent severe infection. This also impacts microbiome acquisition and SCFA levels ^16–18^ and represents another potential avenue by which butyrate availability is reduced in premature infants. Interestingly, alterations in gut microbiome have been linked to increased susceptibility to various diseases of prematurity. Among the SCFAs, butyrate has received considerable attention because of its pleiotropic actions in diverse tissues and pathologic states, including postnatal conditions such as necrotizing enterocolitis, congenital chloride diarrhea, food allergies ^19–25^, and, importantly, ocular diseases such as uveitis ^26, 27^. The purpose of the current study is to test the therapeutic efficacy of butyrate, for the first time ever, in the context of ROP.

## 2. Materials and Methods

### 2.1. Mice

All animal procedures were performed per the statement of the Association for Research in Vision and Ophthalmology (ARVO) and protocols approved by The Augusta University Institutional Animal Care and Use Committee (Approval no: 2009-0214). C57BL/6J mice were purchased from Jackson Laboratories (Bar Harbor, ME) at 2 months of age and were maintained and bred in the Augusta University animal facility (Augusta, GA, USA), with a daily light cycle of 12 hours and fed ad libitum.

### 2.2. Animal Model of Oxygen-Induced Retinopathy (OIR)

The experimental model of ROP, known as Oxygen-Induced Retinopathy (OIR), was implemented in C57BL/6J mice following the protocol outlined by Smith et al. ^28^, which has been previously utilized in our laboratory ^29^. On postnatal day 7 (P7), litters of newborn mice and their respective mothers were placed in a custom-built chamber (the BioSpherix hyperbaric chamber, uniquely designed with iris ports, allowing in vivo treatments in mice without causing a significant decrease in oxygen levels) where the oxygen level was maintained at 75% for 5 days to induce vaso-obliteration. At P12, the mice were returned to room air (RA; 21% oxygen) and kept until P17 to promote retinal neovascularization. To assess the effect on normal retinal development, a separate group of mice was kept in RA from days P0 to P17 as age-matched controls. Butyrate was administered by daily oral gavage (using a specialized cannula designed for the treatment of neonatal mice) at a dose of 200 or 500 mg/kg ^30–32^ or vehicle (PBS; phosphate-buffered saline, 10 μl volume) from postnatal days 7 to 17 in mice undergoing the OIR procedure. Upon completion of the treatment, mice were sacrificed, and eyes were collected for further evaluation.

### 2.3. Hyperglycemia-associated retinopathy (HAR) mice

In this model, C57BL/6J neonatal mice were injected intraperitoneally with a dose of 50 mg/kg bw/day of streptozotocin (STZ) consecutively from P1 to P9 using a 34-G needle (Hamilton syringe), with control animals receiving equal volumes of vehicle (Na-citrate buffer). Under the above conditions, hyperglycemia is induced around P8, leading to delayed retinal vascularization found at P10, as reported previously ^33, 34^. Additional groups were co-injected with STZ and butyrate at a dose of 500 mg/kg, i.p. (10 μL volume) from P1-P9. At P10, retinas were whole-mounted and stained with isolectin B4 (IB4) antibody. Both male and female mice were included, with littermates receiving vehicle (PBS) as a control. The retinal vasculature was analyzed at P10.

### 2.4. Cells and Angiogenesis Assay

Human retinal microvascular endothelial cells (HREC) were obtained from Cell Systems Corporation (Kirkland, WA, USA) and cultured per the manufacturer’s instructions. Cells used in experiments were between passages 3 and 5. The *in vitro* angiogenesis assay was conducted and performed as described previously ^29^. HRECs were treated with vehicle (PBS) or butyrate (1, 2, and 5 mM) for 12 hours under hypoxic conditions (pO2=2%). After treatment, the cells were plated and maintained on Matrigel-coated plates for an additional 6 hours. At the end of this period, the cells were trypsinized, and 300 µL of the cell suspension containing 1.0 x 10^5^ cells/well were added to 24-well plates coated with growth factor-reduced Matrigel matrix (Corning Life Sciences, Tewksbury, MA, USA) according to the manufacturer’s protocol. The angiogenesis assay plate was then incubated at 37°C, 5% CO_2_ for 6 hours. Microscopic images were taken using a Revolve ECHO discover microscope to analyze the vasculature and an unbiased AI-based analysis program (FastTrack AI, iBidi, USA) to calculate total tube length, loop count, and branch count.

### 2.5. Central Vaso-obliteration and Neovascularization

To assess retinal vessel growth and distribution, mice were euthanized at P9 and P17 and eyes were enucleated and fixed with 4% paraformaldehyde for overnight. Retinal flat mounts were prepared and stained with biotinylated Isolectin GS-IB4 from *Griffonia simplicifolia* (0.2 mg/ml; Invitrogen, Carlsbad, CA) and Texas red-conjugated avidin D overnight at 4°C ^29, 35^. Retinal flat mounts were imaged with a Leica Stellaris Confocal microscope. Electronic images were processed using http://oirseg.org ^36^ to create whole retina montages, as previously described ^36^. Areas of vaso-obliteration and neovascularization were automatically calculated.

### 2.6. Western blotting

Total proteins were extracted from the retinas of RA, OIR, and OIR+ NaB mice at P17 days or HRECs using RIPA (Radioimmunoprecipitation assay) cell lysis buffer (Thermo Fisher, Waltham, MA, USA) containing 1% phosphatase and protease inhibitor cocktail (Sigma-Aldrich, St. Louis, MO, USA). An equivalent amount of protein samples (40–60 µg) were subjected to SDS–PAGE and transferred onto a PVDF (Polyvinylidene difluoride) membrane. Then, the membrane was blocked using 5% skimmed milk and incubated with the following anti-mouse primary antibodies: HIF1α (1:1000, Abclonal, Woburn, MA, USA), ICAM1 (1:1000, Abclonal, Woburn, MA, USA), VCAM-1 (1:1000, Abclonal, Woburn, MA, USA), and RBPMS (1:500, Genetex, Alton Pkwy Irvine, CA USA). After immunoblotting, the membranes were stripped using stripping buffer (Boston Bio products, Milford, MA USA) and re-probed with anti-β actin antibody (1:3000; Sigma-Aldrich, St. Louis, MO, USA). The chemiluminescence-based assay was used for band detection (BioRad) using Azure Imaging Systems (Azure Biosystem, Dublin, CA). Scanned images of blots were used to quantify protein expression using NIH ImageJ software (http://rsb.info.nih.gov/ij/). Results were expressed as fold change in protein expression.

Assessment of VEGF_165_ (1:100; Sigma-Aldrich, St. Louis, MO, USA) protein levels was done using heparin affinity columns (Sigma-Aldrich, St. Louis, MO, USA) and Western blot analysis as previously described ^37, 38^.

### 2.7. Immunofluorescent and H&E Staining

The immunostaining of retinal cryosections was performed as described ^29, 39^. Slides were fixed in 4% paraformaldehyde and incubated overnight at 4°C with anti-mouse primary antibodies used at the following concentrations: RBPMS (1:100, GeneTex, Alton Pkwy Irvine, CA, USA), GFAP (1:100; Abcam, Cambridge, MA, USA). Microglia staining was performed on retinal flat mounts using anti-ionized calcium-binding adapter molecule 1 (IBA1; 1:100; FUJIFILM Wako Chemicals Europe; Neuss, Germany) overnight at 4 ◦C. Retinal flat mounts or cryosections were washed three times with 0.1% Triton X-100 in 0.1 M PBS (pH 7.4) followed by incubation with appropriate fluorescence-conjugated secondary antibodies (LifeTechnologies, Eugene, OR, USA). Sections were mounted using fluoroshield mounting medium with DAPI (40,6-diamidino-2-phenylindole; Sigma-Aldrich, St. Louis, MO, USA) and images captured at 20X magnification using a Zeiss LSM 780 Inverted Confocal.

Hematoxylin and eosin (H&E) staining was performed on select P17 retinal and liver samples to assess the toxicity of NaB treatment in these tissues as follows. Samples were enucleated, fixed in 4% paraformaldehyde, sectioned (5 µm), and stained with hematoxylin and H&E as per previous protocols ^39^.

### 2.8. Statistical Analysis

The results are presented as mean ± SD for 3-6 replicates. The potential significance of differences between the treatment groups was evaluated using one-way ANOVA followed by Dunnett’s multiple-comparison test, and results were considered significant at *p* < 0.05. The graphs were prepared using GraphPad Prism version 10 software (GraphPad, San Diego, CA, USA).

## 3. Results

### 3.1. Effects of oral butyrate administration on retinal vasoobliteration and neovascularization in OIR mice

The OIR mouse model is a well-established rodent model in which to study key pathologic characteristics relevant to human ROP. Using this model, we evaluated the effects of butyrate on retinal neovascularization. A schematic overview of the OIR mouse model and treatment protocol is presented in Fig. 1A. In brief, OIR mice were treated with sodium butyrate (NaB; 200 and 500 mg/kg p.o.) or vehicle (PBS) by daily oral gavage from postnatal days 7 to 17. All mice were euthanized at P17, and retinal flat mounts were prepared and stained with isolectin B4 to highlight the retinal vasculature. As shown in Fig. 1B, subtle, potentially non-significant improvements were evident in OIR retinas treated with 200 mg/kg NaB. However, at the higher dose of 500 mg/kg NaB treatment, there was a significant reduction in retinal vessel loss and in neovascularization (Figs. 1D-E), contributing overall to a more uniform vascular network. This is perhaps better visualized in Figs. 1F-H, in which the same retinal flat mount images are shown in grayscale with areas of neovascularization and vasoobliteration areas highlighted in red and yellow, respectively. To verify these qualitative findings, we used oirseg, an unbiased automated web application, to quantitatively assess differences in vaso-obliteration (Fig. 1I) and neovascularization (Fig. 1J) in retinal flat mounts prepared from study mice. Morphometric analysis showed that butyrate (500 mg/kg) significantly decreased avascular area and neovascularization when compared to OIR mice (Fig. 1I-J).

**Figure 1:**
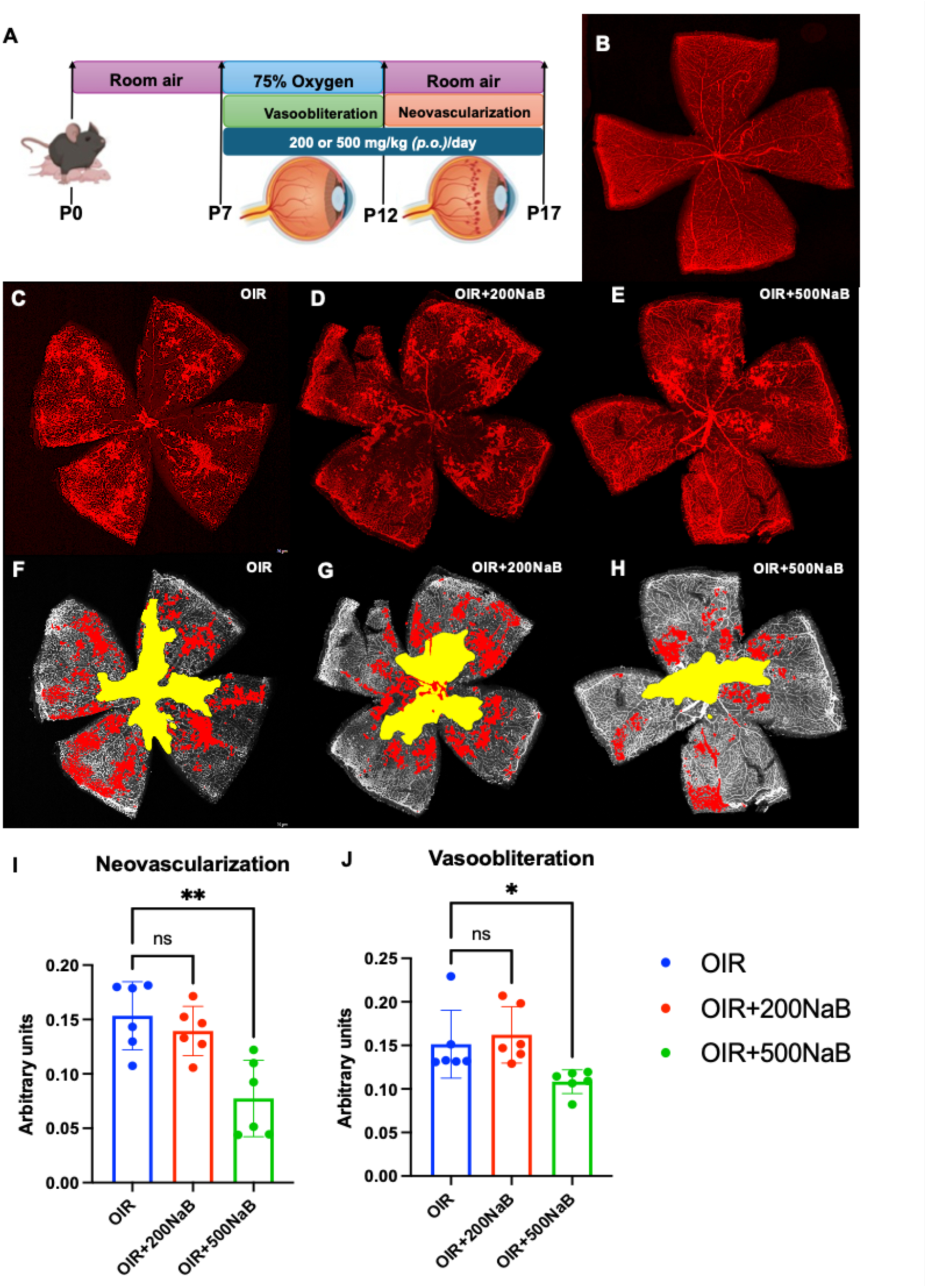
Effects of butyrate administration on retinal vaso-obliteration and neovascularization in OIR mice. Butyrate (200 or 500 mg/kg) or vehicle (PBS) was administered daily by oral gavage from postnatal days 7 to 17. All mice were euthanized at P17, and retinal flat mounts were stained with isolectin B4. **(A)** Schematic representation of the mouse model of oxygen-induced retinopathy (OIR). (**B-H)** Representative retinal flat mounts from experimental groups. Histograms represent the morphometric analysis of retinal flat mounts measuring **(I)** vaso-obliteration and **(J)** neovascularization in OIR mice calculated using an unbiased automated web application (oirseg). Values are mean ± SD (n=6). *p<0.05 vs. OIR (Vehicle-treated).

### 3.2. Effect of butyrate on in vitro tube formation in hypoxic human retinal microvascular endothelial cells (HREC)

Building upon our *in vivo* vascular findings of increased neovascularization in OIR mice, a feature that was reduced in association with butyrate treatment, we examined the impact of butyrate on retinal endothelial cells, employing an *in vitro* model of hypoxia-induced tube formation in HRECs. HRECs were cultured at 37°C in a humidified atmosphere of 5% CO_2_ as per the manufacturer. Cells were exposed to 12 hours hypoxia (2% oxygen), treated with different concentrations of butyrate (1, 2, and 5 mM) or kept untreated (normoxia) for 12 hours, and then plated onto a Matrigel matrix to induce tube formation *in vitro*. As shown in Fig. 2, the formation of tube-like structures, a process relevant to endothelial cell angiogenic function, was significantly induced in hypoxic conditions (Fig. 2A). Tube formation was dose-dependently inhibited by butyrate treatment. Indeed, at the doses of 2 and 5 mM (Figs. 2C and 2D, respectively), butyrate significantly decreased total tube length and number of branch points and loop count compared to vehicle (PBS) treated hypoxic endothelial cells (Fig. 2F-H). These *in vitro* findings agreeably align with our *in vivo* results, reinforcing the direct and promising impact of butyrate on retinal endothelial cell function and its therapeutic potential in OIR/ROP.

**Figure 2:**
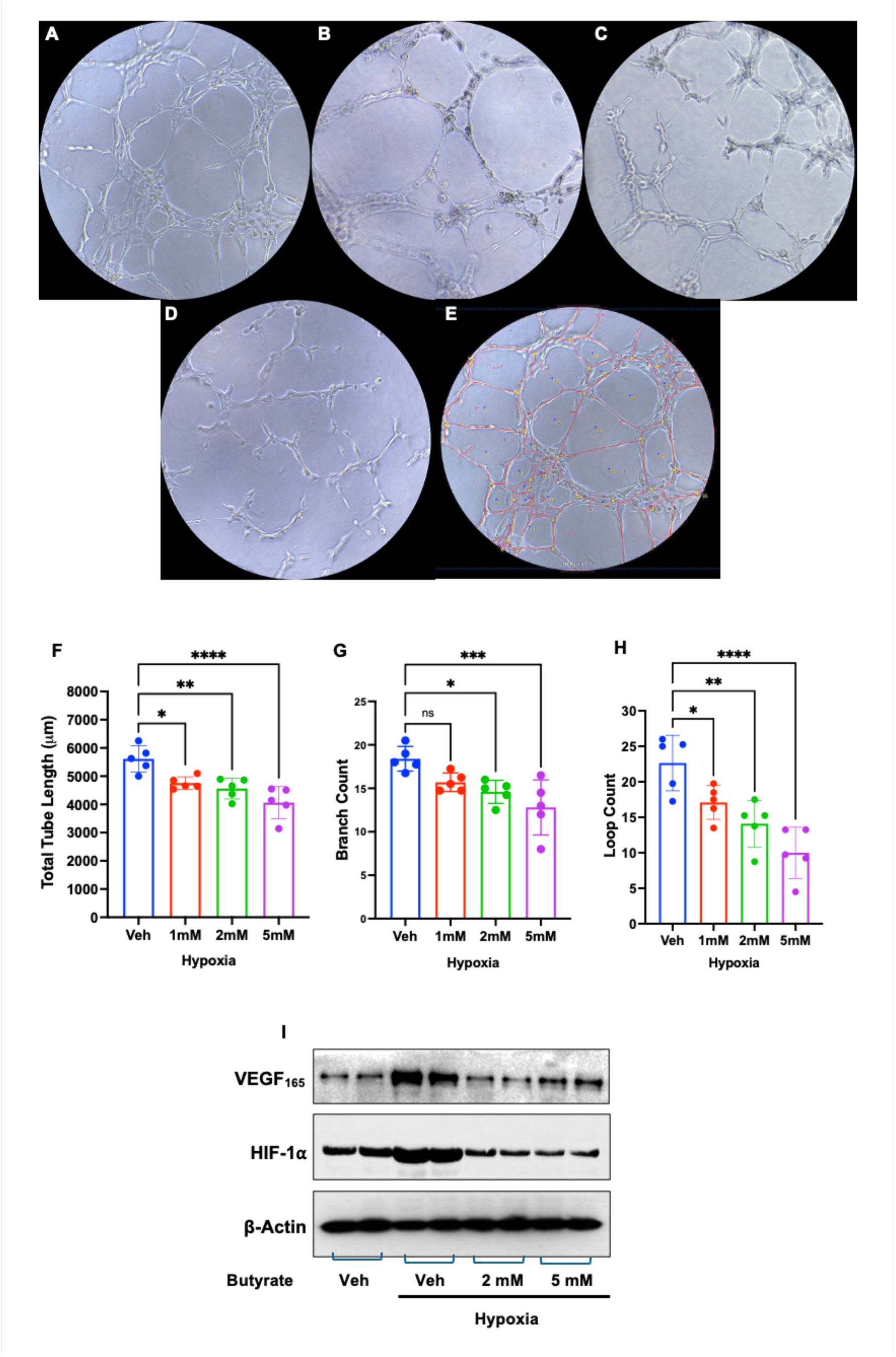
Effect of butyrate on *in vitro* tube formation in hypoxic human retinal microvascular endothelial cells (HRECs). HRECs were treated with vehicle (PBS) or butyrate (1, 2, and 5 mM) for 12 hours under hypoxic conditions (pO_2_=2%) and later plated and maintained on Matrigel-coated plates for an additional 6 hours. **(A-E)** Microscopic images were taken and used for the analysis of vasculature using an unbiased AI-based analysis (FastTrack AI, iBidi, USA) to calculate **(F)** total tube length, **(G)** loop count, and **(H)** branch count. **(E)** A representative image processed via the analysis software shows how each parameter was obtained. Values are mean ± SD (n=5; with four experimental replicates in each group). *p<0.05 vs. Vehicle-treated cells. **(I)** Western blot analysis was conducted on cell lysate from each group to measure VEGF_165_ and HIF-1α levels, normalized to β-actin.

Hypoxia-inducible factor-1 (HIF-1α) is a master transcriptional regulator of cellular responses to hypoxia and plays a key role in retinal neovascularization ^40–42^. We additionally evaluated the expression of HIF-1α and its downstream angiogenic mediator, VEGF protein expression, in HRECs cultured in normoxic or hypoxic conditions in the presence or absence (vehicle control) of butyrate. As shown in Fig. 2I, butyrate treatment decreases HIF-1α protein expression substantially in HREC exposed to hypoxia, resulting in a significant decrease in hypoxia-induced VEGF_165_ expression. As such, it is plausible that butyrate regulates HIF-1α at transcription or post-transcriptional levels to influence downstream angiogenic mediators such as VEGF_165_.

### 3.3. Effect of butyrate administration on retinal ganglion cell survival and reactive gliosis in OIR mice

The retina is composed of different cell types in addition to endothelial cells. Hematoxylin and eosin (H&E)-stained retinal cross-sections (Figs. 3A-C) were prepared such that the potential impact of OIR and butyrate on the neuroretina of mice could be evaluated. The structural architecture of the neuroretina remained largely intact in OIR mice in the presence and absence of butyrate treatment. However, large clusters of vessel tufts could be detected in the inner retina at the vitreoretinal interface in OIR mice. The size and number of vascular tufts were reduced in OIR mice treated with butyrate (Figs. 3D). This aligns well with the findings described in Figs. 1-2 above.

**Figure 3:**
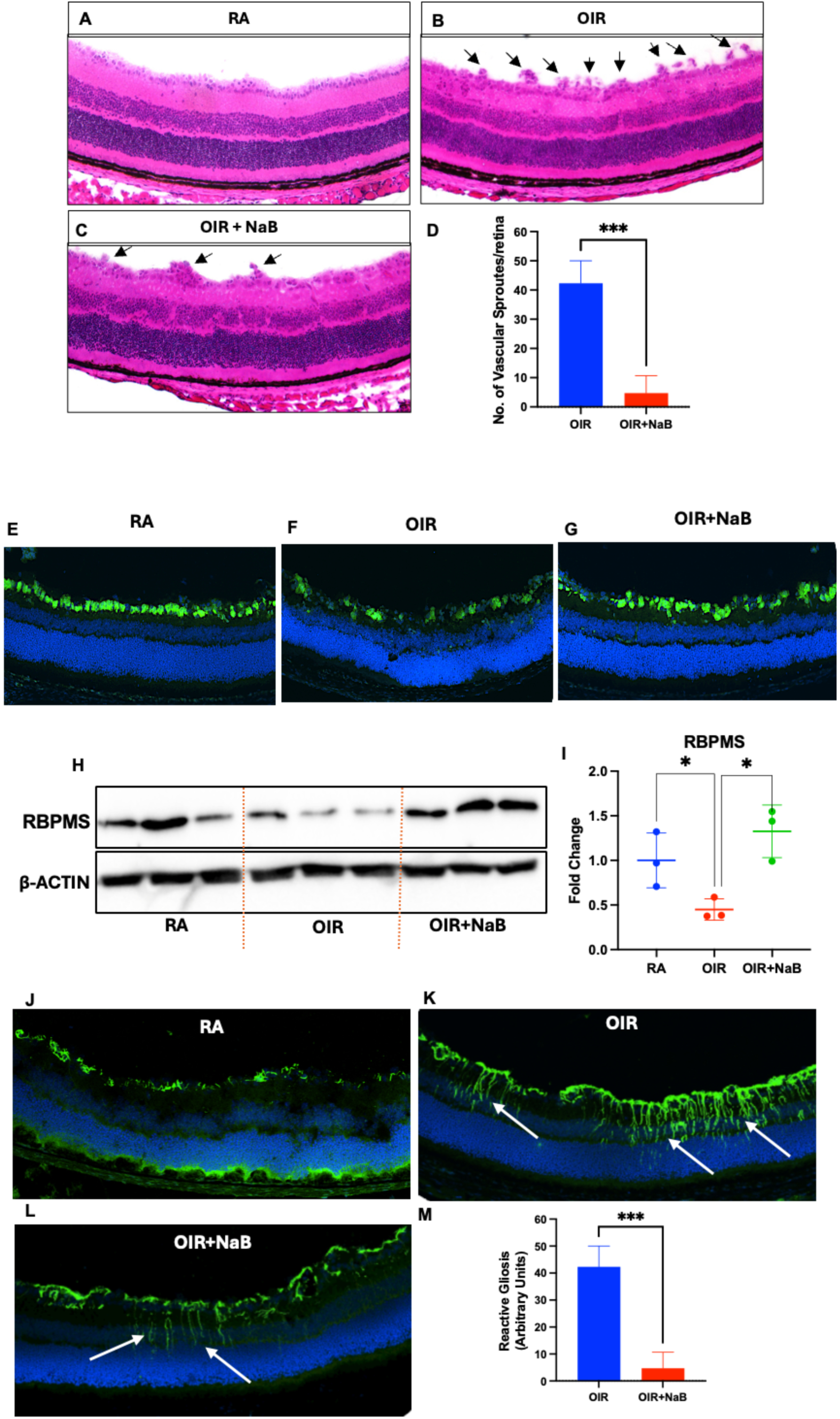
Effect of butyrate administration on retinal histoarchitecture, retinal ganglion cell survival, and reactive gliosis in OIR mice. Light micrograph of retinal cross-sections from P17 mice, stained with H&E at 20X magnification. Images show the retina from a **(A)** RA, **(B)** OIR, and **(C)** OIR+NaB. Arrows indicate the neo-vessels. **(D)** Neovessels were counted within 200 μm from either side of the optic nerve at 20X. Immunostaining was performed on samples from room air (RA), oxygen-induced retinopathy (OIR), and OIR treated with Butyrate (NaB, 500 mg/kg) from P7 to P17 (OIR+NaB) in mice using **(E-G)** RBPMS (RNA-binding protein with multiple splicing is a protein) to identify retinal ganglion cells, and **(J-L)** GFAP (Glial fibrillary acidic protein) to identify retinal Müller cells activity (reactive gliosis). **(M)** Reactive gliosis was quantified by counting the number of GFAP-positive inter-retinal projections (marked by arrow). **(H-I)** Western blot analysis was conducted using retinal protein extracts from each group to measure RBPMS, normalized to β-actin. The bar graphs present the quantitative analysis of 3-4 independent mouse retina samples (n=3), expressed as mean ± SD. *p<0.05 compared to the OIR (vehicle-treated) group.

Inflammation is a key factor in the development and progression of human ROP. This is also true in the OIR model. The retina is particularly susceptible to pro-inflammatory insults. Reactive gliosis and loss of retinal ganglion cells are features associated with inflammation and OIR in mice ^43^. Although H&E-stained retinal cross-sections (Fig. 3) appeared to be grossly normal with the exception of the changes noted above, the location of the vascular anomalies in the inner retina may mask potential changes that impact the ganglion cell layer. To assess the effects of butyrate on retinal ganglion cells in OIR mice, we performed immunofluorescence and Western blot analyses using the ganglion cell-selective marker RNA binding protein with multiple splicing (RBPMS) ^44^. The results of these analyses showed a marked decrease in RBPMS-specific immunoreactivity in OIR mice retinas that was limited in animals that received butyrate treatment (Fig. 3E-I). Further, we observed significant reactive gliosis in OIR mice, indicated by increased glial fibrillary acidic protein (GFAP) immunoreactivity throughout OIR mouse retinas. Reactive gliosis was significantly reduced in the retinas of OIR mice treated with butyrate (Fig. 3 J-M).

### 3.4. Effect of butyrate administration on microglial activation in OIR mice

Microglia, the resident macrophages responsible for maintaining retinal integrity, play a crucial role in regulating immunity and inflammatory responses in the retina. There is a burgeoning literature suggesting that these cells may play a key role in the development and progression of ROP. Microglia display distinct phenotypical changes in active and inactive states ^45^. Immunofluorescence localization of ionized calcium-binding adaptor molecule 1 (Iba-1), a macrophage and microglial selective marker, was performed on retinal flat mounts to assess microglia polarization in OIR mouse retinas in the presence and absence of butyrate treatment (Fig. 4A-D). Ramified (quiescent) microglia were predominant in retinal flat mounts prepared from RA control mice. However, OIR flat mounts were riddled with amoeboid (activated) microglia. In line with our observations above, NaB treatment significantly reduced microglia activation, a sign of reduced inflammation, in OIR, as evidenced by a reduction in the numbers of amoeboid-shaped microglial cells in OIR+NaB retinal flat mounts (Fig. 4A-D). To follow up on this finding, we monitored the expression of intracellular cell adhesion molecule (ICAM-1) and vascular cell adhesion molecule (VCAM-1), adhesion molecules considered to be primary drivers of neovascularization in diabetic retinopathy ^46^ and other degenerative diseases in which pathological neovascularization is common ^47^. VCAM-1 and ICAM-1 expression were induced in the OIR retinas at P17 as compared to RA mice (Fig. 4E-G). However, butyrate treatment was associated with a decrease in the expression of these pro-inflammatory mediators in OIR conditions.

**Figure 4:**
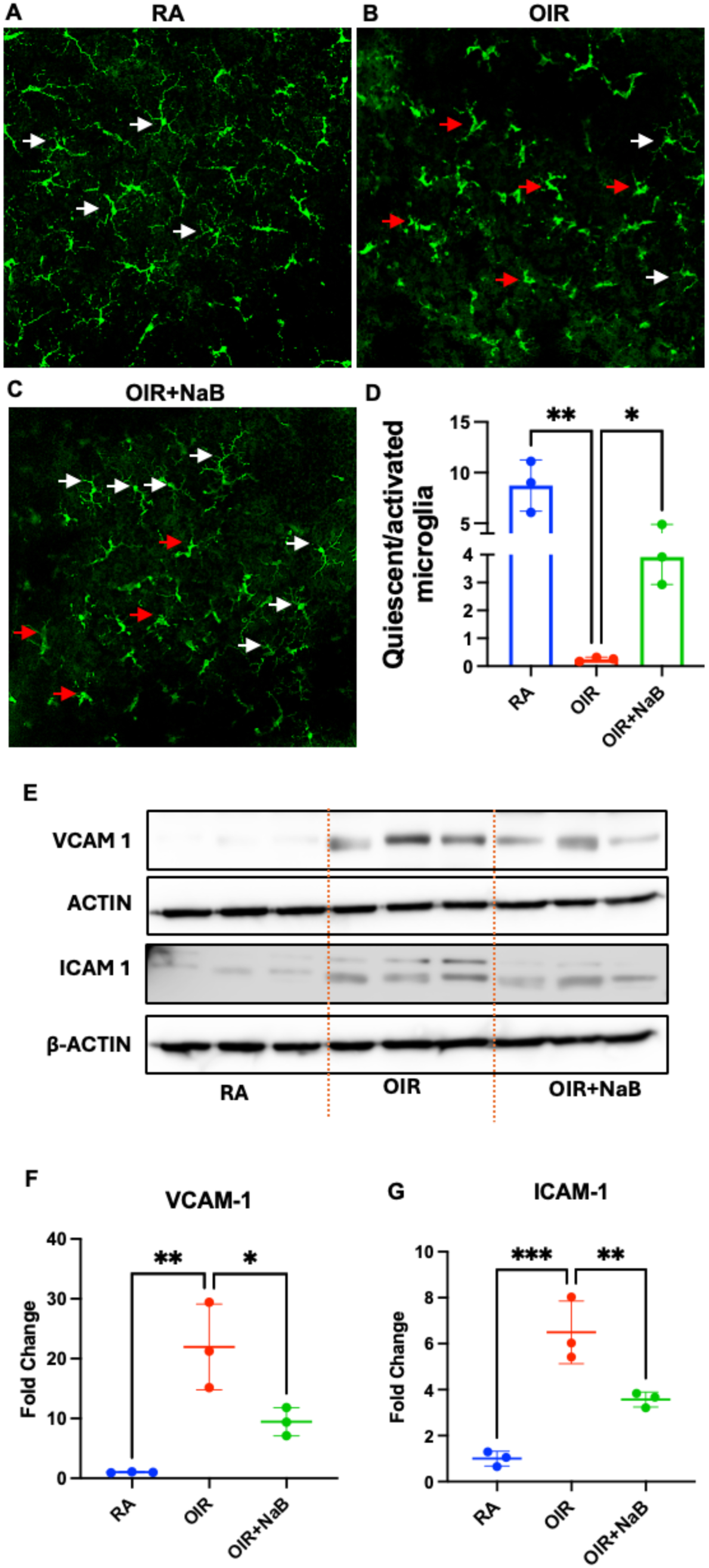
Effect of butyrate administration on microglial activation in OIR mice. Immunostaining was performed on samples from room air (RA), oxygen-induced retinopathy (OIR), and OIR treated with Butyrate (NaB, 500 mg/kg) from P7 to P17 (OIR+NaB) in mice. **(A-D)** Retinal flat mounts were immunostained for microglia using Iba-1, showing quiescent (white arrows) and activated states (red arrows) from n=4-stained retinal sections for each group. **(E)** Western blot analysis was conducted on the retinal protein extracted from each group to measure VCAM-1 and ICAM-1 levels, normalized to β-actin**. (F-G)** The bar graph presents the quantitative analysis of 3 independent mouse retina samples (n=3), expressed as mean ± SD. *p<0.05 compared to the OIR (vehicle-treated) group.

### 3.5 Effect of butyrate administration on high-oxygen and hyperglycemia-associated retinal vaso-obliteration in mice

Current clinical management of ROP focuses on phase II of the disease. There are no therapies to impact early (phase I) ROP. To determine whether oral butyrate supplementation could be one such potential therapy, we monitored OIR at P9 (Fig. 5A-C), a time-point consistent with early ROP in humans, as opposed to P17, as done in our prior experiments. Indeed, butyrate was also effective at reducing vasoobliteration in this early phase (phase I) of OIR.

**Figure 5:**
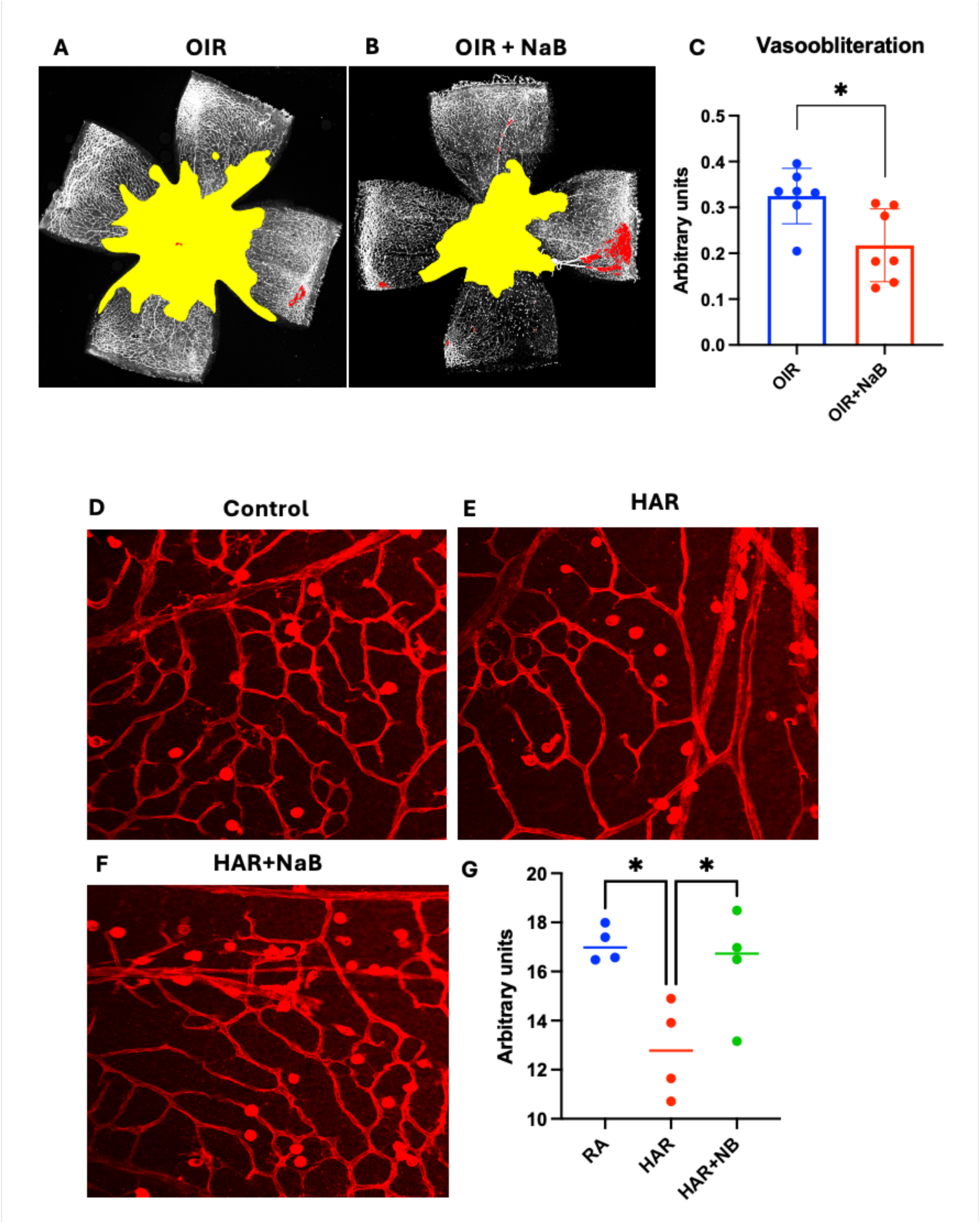
Effect of butyrate administration on high-oxygen and hyperglycemia-associated retinal vasoobliteration in mice. **(A-B)** Representative images of Isolectin-B4-stained retinal flat mounts from vehicle and butyrate (NaB, 500 mg/kg) treated OIR mice at P9. **(C)** Histograms display the morphometric analysis of retinal flat mounts, measuring vaso-obliteration calculated with the unbiased automated web application (oirseg). Data are presented as mean ± SD (n=6). *p<0.05 compared to the OIR (vehicle-treated) group. **(D-F)** Representative images of Isolectin-B4-stained retinal flat mounts (20X) from control, HAR, and HAR treated with 500 mg/kg butyrate (HAR+NaB) mice. **(G)** Vascular density was calculated using the ImageJ vessel analysis plug-in. Data are presented as mean ± SD (n=4). *p<0.05 compared to the HAR (vehicle-treated) group.

Because of the long-known adverse effects of variations in oxygen concentrations on the developing retina, oxygen concentrations are now tightly regulated in most neonatal intensive care units; however, ROP persists, making it clear that factors other than oxygen must be involved. Hyperglycemia is also common in premature infants and is considered a significant risk factor for ROP ^34^. Using the recently developed HAR mouse model ^34^, a model that recapitulates key early pathologic features in the retina consistent with phase I ROP, we evaluated the impact of butyrate therapy. Mice were injected daily with streptozotocin (STZ, 50 mg/kg) into the peritoneal cavity from postnatal day 1 (P1) to postnatal day 9 (P9) to induce hyperglycemia. Mice were then treated with either vehicle (PBS) or 500 mg/kg butyrate (NaB+ HAR). Non-STZ-injected mice served as controls. On postnatal day 10 (P10), body weight and blood glucose levels (Data not shown) were recorded, and retinas were harvested. Retinal flat mounts were stained with isolectin B4 to assess vascular distribution. As shown in Fig. 5D-G, butyrate treatment resulted in a significantly reduced avascular area compared to HAR mice (Fig. 5E). Further, the formation of the superficial vascular network was notably improved in NaB+ HAR mice relative to HAR mice (Fig. 5D-G), indicating enhanced retinal neovascular maturation in the butyrate-treated group.

## 4. Discussion and Conclusions

ROP is a leading cause of blindness in children worldwide ^1–3^. Despite improvements in neonatal care, the disease remains a significant threat to the vision of children born prematurely. Current clinical management of the disease involves surgical and therapeutic strategies that focus on phase II of the disease and are thereby of use relatively late, after much damage to the retina, although potentially undetectable by routine clinical examination, has already occurred. Further, although these strategies may be effective in limiting the abnormal neovascular component of the disease in some premature infants, many of them continue to develop significant visual problems as they progress into childhood ^4–6^. As such, new, improved therapies for preventing and treating ROP represent a major unmet clinical need.

The current study focuses on understanding better the roles of vascular and non-vascular cells in the retina as it relates to the pathogenesis of ROP. Importantly, we also tested a novel orally delivered therapy, NaB, in preclinical *in vivo* and *in vitro* models of the disease. Our findings demonstrate that daily oral supplementation of the SCFA provides significant protection against pathologic angiogenesis in the oxygen-induced retinopathy (OIR) mouse model, a model used widely to evaluate key environmental and pathologic features consistent with ROP development in human infants. Further, we show that not only does butyrate impact vascular pathology but also pathology affecting neuronal and microglial cells in the inner retina, cell types that have been considerably understudied compared to endothelial cells in OIR/ROP. Our early evaluations in P17 OIR mice demonstrate that butyrate may be an effective alternative or adjuvant therapy to anti-VEGF or other current strategies to impact ROP. Importantly, studies in OIR mice at P9 and in the novel HAR mouse model, a model that recapitulates the hyperglycemic environment present in many premature infants that ultimately develop ROP ^34^, show butyrate to be an effective therapy for intervening much earlier in the sequence of events, potentially during phase I, that lead to clinically detectable retinal pathology in ROP.

The current study represents the first, to our knowledge, to investigate the potential of butyrate as a therapy to prevent and treat ROP. Further, it provides insight into the impact of butyrate therapy on multiple retinal cell types (e.g., endothelial, microglial, and ganglion) and mechanisms (e.g., hypoxia/hyperoxia, inflammation, hyperglycemia) relevant to early (phase I) and advanced (phase II) ROP. Though additional studies are warranted to further decipher the specific mechanistic pathways by which butyrate directly or indirectly impacts the development and progression of ROP-like retinal pathology in OIR and HAR mice, the current study sets the stage for additional preclinical testing and optimization of butyrate in ROP. Indeed, the fact that butyrate impacts OIR/ROP at multiple stages increases the translational appeal of the current findings, with the present study setting the stage for further detailed investigations.

## Funding

We want to acknowledge funding support for these studies from Augusta University Start-up funds to RNJ. The corresponding author’s laboratory work is funded by EY035336 and the MMC-CZI accelerated precision health program (RNJ).

## Authors’ Contributions

Conceptualization, P.M.M. and R.N.J.; Data analysis and curation, A.O., T.B., J.L., G.S., and A.J.B; Conducted experiments, A.O., T.B., J.L., G.S., and A.J.B; Resources, M.C.T, P.M.M and R.N.J; Writing – original draft, M.C.T, P.M.M. and R.N.J.; Validation, A.O., T.B., J.L., Review-writing, M.C.T, P.M.M and R.N.J. All authors contributed to the article and approved the submitted version.

## Acknowledgment

We (T.B.) acknowledge support in part by the Augusta University Provost’s Office and the Translational Research Program of the Department of Medicine, Medical College of Georgia at Augusta University.

## Availability of data and materials

The datasets supporting the conclusions of this article are included within the article; further inquiries can be directed to the corresponding author.

## Declaration of competing interest

The author(s) declared no potential conflicts of interest with respect to the research, authorship, and/or publication of this article.

## Generative AI statement

The authors declare that no Generative AI was used in the creation of this manuscript.

## Notes

### Competing Interest Statement

The authors have declared no competing interest.

### Summary of Updates

Changes in Author affiliations, new data and additional analysis

